# MukBEF-dependent chromosomal organization in widened *Escherichia coli*

**DOI:** 10.1101/2022.07.13.499882

**Authors:** Aleksandre Japaridze, Raman van Wee, Christos Gogou, Jacob W. J. Kerssemakers, Cees Dekker

## Abstract

The bacterial chromosome is spatially organized through protein-mediated compaction, supercoiling, and cell-boundary confinement. Structural Maintenance of Chromosomes (SMC) complexes are a major class of chromosome-organizing proteins present throughout all domains of life. Here, we study the role of the *Escherichia coli* SMC complex MukBEF in chromosome architecture and segregation. Using quantitative live-cell imaging of shape-manipulated cells, we show that MukBEF is crucial to preserve the toroidal topology of the *E. coli* chromosome and that it is non-uniformly distributed along the chromosome: it prefers locations towards the origin and away from the terminus of replication, and it is unevenly distributed over the origin of replication along the two chromosome arms. Using an ATP hydrolysis-deficient MukB mutant, we find that MukBEF translocation along the chromosome is ATP-dependent, in contrast to its loading onto DNA. MukBEF and MatP are furthermore found to be essential for sister chromosome decatenation. We propose a model that explains how MukBEF, MatP, and their interacting partners organize the chromosome and contribute to sister segregation and recombination. The combination of bacterial cell-shape modification and quantitative fluorescence microscopy paves way to investigating chromosome-organization factors *in vivo*.

## Introduction

The intricate organization of genetic material in chromosomes remains incompletely understood, even in thoroughly studied bacteria such as *Escherichia coli* (*E. coli*). In order to fit into the volume of a single cell, the nucleoid needs to be strongly compacted^1^, whilst preserving a complex spatial and dynamic organization that facilitates vital cellular processes. In *E. coli*, compaction is achieved by a combined interplay of DNA supercoiling^2^, nucleoid associated proteins (NAPs^3^), and a Structural Maintenance of Chromosomes (SMC) complex called MukBEF^4–9^. MukBEF is a pentamer consisting of double copies of MukB and MukE subunits and a single kleisin unit called MukF^9^. DNA can be bound and re-shaped *in vitro* by the hinge-like configuration of the MukB subunits in an ATP-dependent manner^10^. Recent studies reported that a 6-fold increase in the number of MukBEF copies led to the formation of a ring-like structure of SMCs along the toroidal nucleoid^11,12^. This ring was hypothesized to function as a chromosomal backbone from which peripheral DNA loops protrude.

MukBEF also mediates chromosomal interactions with other proteins that organize and disentangle sister chromosomes during replication and segregation^13^. One such protein is the so-called Macrodomain ter Protein (MatP). MatP’s organizational role is commonly associated with its active displacement of MukBEF from the ter region^13^. MatP binds *matS* sites near the terminus of replication and localizes the ter macrodomain to the midcell through a direct interaction with the divisome^14^. There is growing *in vivo* evidence that in the absence of MatP, MukBEF is unable to be displaced from the ter region which then results in severe condensation of the ter region^11,13,15^. Recently, the molecular structure of the MukBEF-MatP-*matS* nucleoprotein complex was resolved using cryo-EM, revealing how the subunits of MukBEF and MatP directly interact^16^. *matS*-bound MatP was found to sit at the center of the MukBEF ring, potentially blocking MukBEF translocation in the ter domain and promoting ATP-dependent un-loading of the SMC *in vivo*. Furthermore, deletion of MukBEF was shown to result in anucleation and defects in chromosome segregation^17–19^, whereas deletion of *matP* led to premature segregation of sister foci in the ter macrodomain and their mis-localization relative to the divisome before cell division^13,15^.

Another important direct interaction partner of both MukBEF and MatP is the topoisomerase IV^20–22^. Topo IV influences the linking number of the nucleoid primarily in the ori region, where MukBEF is predominantly localized. Furthermore, it has been found to mediate the timely segregation throughout replication as well as the decatenation of sister chromosomes after replication^23–25^. Although these findings form a foundation for the understanding of chromosomal organization, the high degree of nucleoid compaction in combination with simultaneous ongoing replication cycles have so far impeded direct *in vivo* visualization of their actions.

To map spatiotemporal localizations and investigate the interactions of MukBEF and MatP with the chromosome, we employed a method to synchronize chromosome replication in a population of *E. coli* cells (by using a temperature-sensitive *dnaC* allele^26^) and simultaneously increased their size through cell-shape manipulation. For the latter, treatment of cells with low doses of the A22 inhibited the polymerization of the MreB filaments, thereby disrupting the typical rod shape of *E. coli*^27^. These cells gradually expanded in size and typically reached at least 2-fold larger width and length. Concomitantly, the spatial constraint that the cell wall imposed on the nucleoid was thus reduced. In previous studies, we showed how the chromosome in such expanded shapes exhibited a toroidal topology and remained physiologically active in the cell, preserving its capability to replicate and segregate its chromosomes, suggesting that treatment with A22 did not impact cell viability^28–30^. Interestingly, similar widened cell wall-deficient bacteria^31^ were also observed in patients with recurring infections, suggesting that bacterial cells are capable of naturally re-shaping their size and cell wall composition.

Here, we use quantitative fluorescence imaging to study the distribution of MukBEF along the chromosome and we characterize structural changes that result from the mutation or deletion of MukBEF subunits. We show that MukBEF positions along the chromosome with a strong ori-proximal and ter-distal spatial bias in the presence of MatP near the terminus of replication. The preferential localization of MukBEF away from ter is strongly dependent on its ability to hydrolyze ATP, as is its ability to compact the nucleoid. Additionally, MukB is found to spread asymmetrically over the origin of replication along the chromosome arms. We show that the deletion of *matP* does not alter the capacity of MukBEF to bind and compact the nucleoid, but that its localization along the genome is directed by MatP. Upon deletion of *matP*, MukBEF displays a 3-fold increased presence near the ter region which leads to a local compaction of this domain, and which results in severe segregation defects. Deletion of either MukBEF or MatP was found to impair sister decatenation and recombination, resulting in the formation of dimer chromosomes. Our quantitative fluorescence analysis in combination with increased spatial resolution in live shape-modified cells offers new means for investigating chromosome organization *in vivo*.

## Results

### MukBEF spreads non-uniformly along the chromosome

Throughout this study we performed simultaneous four-color imaging of the chromosome (DAPI or HU-mYpet) and MukB (mYpet) together with the origin and the terminus of replication (via Fluorescence Repressor Operator Systems^32^) (Fig.1a). Rod-shaped cells grown in minimal media typically display one or two origins of replication and a single terminus, indicating that cells are in the process of replication^33^ (Fig. 1b). In order to circumvent the optical limitations due to high degree of chromosome compaction and cell-to-cell variability due to ongoing replication cycles, we performed experiments with A22-widened temperature-sensitive (*dnaCts*) *E. coli* cells^26^. Cells were first synchronized by growing above permissive temperature (and hence cells maintained only a single chromosome^26,34^) and were then incubated withA22 to grow larger in size.

**Figure 1.**
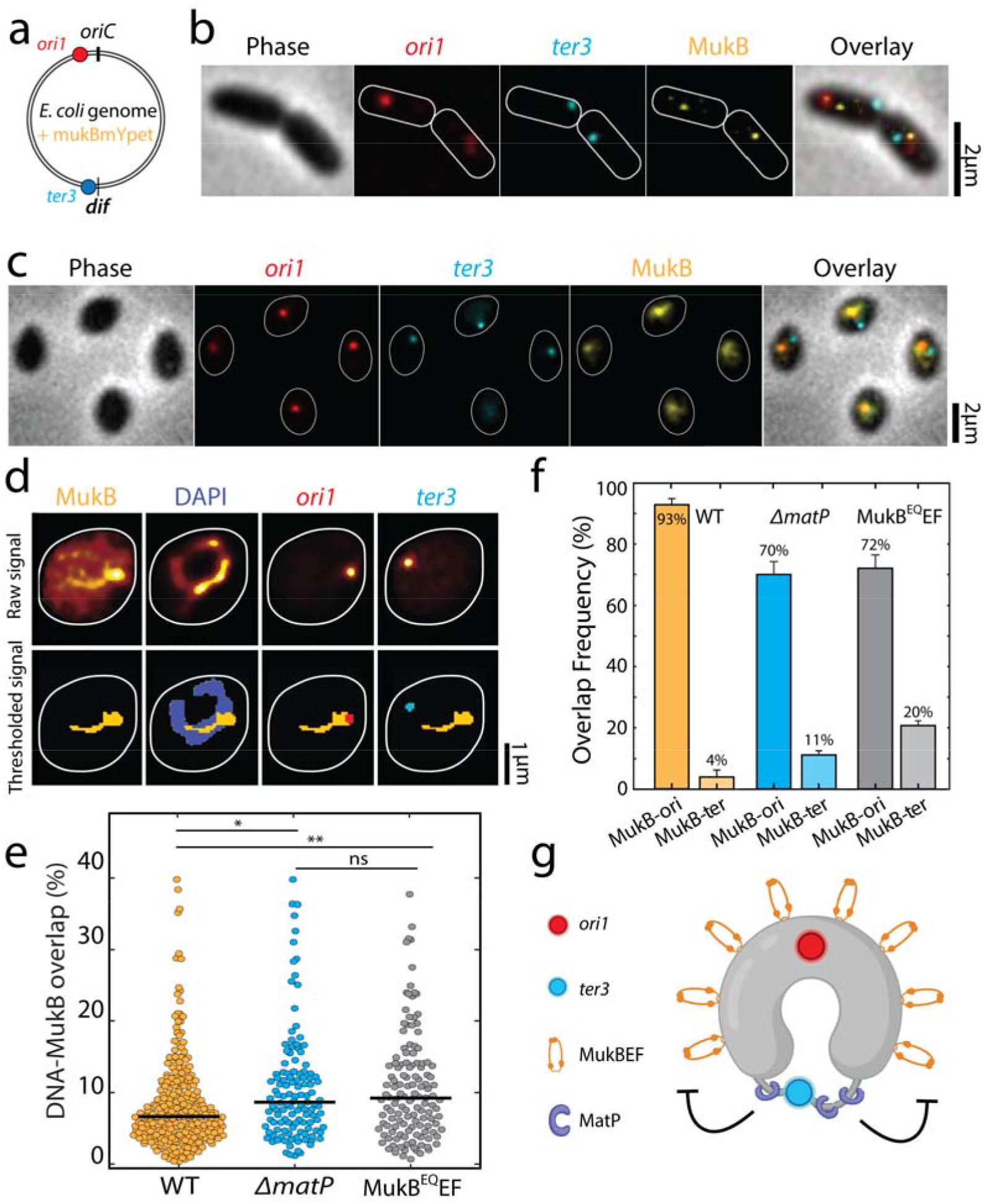
Quantitative localization of MukB complexes along the chromosome. **a**. Schematic representation of a circular *E. coli* genome with FROS arrays at *ori1* and *ter3* location and MukB-mYpet labeling. **b**. Representative image of rod-shaped and **c**. widened *E. coli* cells in phase contrast and the fluorescence channels of *ori1, ter3* and MukB. Cell outline is indicated in white. **d**. Typical example of the thresholding process to calculate fluorescence signal overlap. **e**. Percentage of MukB intensity that overlaps with the DNA mask for wildtype (*N=260 cells*), ⊿*matP* (*N=121 cells*), and MukB^EQ^EF mutants (*N=131 cells*). f. Signal overlap between MukB and *ori1* and MukB and *ter3* foci in wildtype (*N=260 cells*), ⊿*matP* cells (*N=121 cells*) and MukB^EQ^EF cells (*N=131 cells*). Error bars report on the standard deviation. **g**. Schematic representation of MukBEF positioning along the circular *E. coli* chromosome and its regulation by MatP. Statistical significance was determined by performing a single factor ANOVA test. The following conventions are used: ns: 0.05 < p, *: 0.01 < p < 0.05, **: 0.001 < p < 0.01. We report a significant difference in results if p < 0.05.

In these widened cells, the chromosomes organize into a toroidal configuration, with the origin and terminus of replication positioned at opposite halves of the chromosome ring^28,29^. Surprisingly, in the widened cells the MukBEF complexes did not form a single tight cluster as they do in rod-shaped cells (Fig. 1c). Rather, the MukBEF signal distributed over the toroidal chromosome to adopt a significantly extended cluster with a Ferret diameter of 0.6 ± 0.3 µm (mean ± sd, *N=235 cells*) compared to a diffraction-limited diameter of 0.4 ± 0.1 µm (mean ± sd, *N=118 cells*) for the clusters in rod-shaped cells (****: p < 0.0001, Fig. S1). In the widened cells, the signal spread along the left and right arms of the chromosome and the number of MukBEF clusters was typically one or two, as was the number of ori foci in a cell (Fig. 1d).

Next, we measured the ability of MukBEF to bind the nucleoid. We determined the percentage of the nucleoid that was covered by MukBEF clusters (Fig. S2). In wildtype cells, we found that 8.6 ± 0.5 % (mean ± sd, *N=260 cells*) of the total DNA material was co-localized with the MukBEF (Fig. 1e). We compared this with two mutants: ⊿*matP* cells, in which the MatP protein was deleted, and MukB^EQ^EF cells, whose MukB subunits were impaired in hydrolyzing ATP^35^. For the ⊿*matP* and MukB^EQ^EF mutants, this overlap increased to 10.8 ± 0.8 % (mean ± sd, *N=121 cells*) and 10.8 ± 0.9 % (mean ± sd, *N=131 cells*), respectively. We hypothesize that this enhanced coverage of the DNA by MukBEF is because MukBEF also partly occupies the ter region in the mutants, indicating that both the MatP protein and MukB ATPase activity are required for MukBEF displacement.

Furthermore, we quantitatively evaluated MukBEF positioning along the chromosome, by investigating the co-localization of MukBEF with the origin and terminus of replication (Fig. 1f). In 93 ± 2 % of all cells (mean ± sd, *N=260 cells*), MukB and ori signal overlapped, whereas we only observed a 3.8 ± 2.0 % (mean ± sd, *N=260 cells*), overlap between MukB and the ter signal (Fig. 1f). In the remaining 2% of cells, MukB did not colocalize with either of the foci. In ⊿*matP* cells, we found that MukBEF co-localization with the ori was preserved, but to a clearly lesser extent than in wildtype cells (Fig. 1f): in only 70 ± 5 % (mean ± sd, *N=121 cells*) of ⊿*matP* cells, the MukBEF signal was overlapping with ori. This reduction in co-localization could be explained by a redistribution of MukBEF towards the MatP-depleted ter region. Indeed, we observed a 3-fold increase in co-localization of MukB with the ter in ⊿*matP* cells to 11 ± 1 % (4% in wildtype cells). Furthermore, the fraction of cells in which MukB did not overlap with either focus was 10-fold higher than in wildtype (20% vs. 2%). MatP thus is crucial to displace MukBEF from the ter macrodomain. In wildtype cells, MatP is observed as spots flanking the stretched-out ter region (Fig. S3). Deletion of *matP* leads to compaction of the ter region, and subsequent reorganization of the chromosome (Fig. S4). The MukB distribution along the nucleoid of MukB^EQ^EF mutants was similar to the ⊿*matP* mutants, as MukB overlapped with the ori focus in 72% of MukB^EQ^EF cells (*N=131 cells*), 20% of the cells showed an overlap with the ter focus, and the remaining 8% showed no overlap with either. The 5-fold increase (20% vs. 4%) in co-localization of the MukB signal with the ter region for the MukB^EQ^EF cells relative to the wildtype is remarkable as it shows that MukBEF can bind to the ter region but apparently is unable to get displaced if its capability to hydrolyze ATP is impaired, in agreement with earlier work^13^.

### MukBEF is distributed asymmetrically over the left and right chromosome arms

Visualizing the toroidal structure of the chromosome and separately observing the two arms of the chromosome enabled us to zoom in further on the spreading of the MukB in wildtype cells, and ⊿*matP* and MukB^EQ^EF mutants (Fig. 2ab). We first measured the relative local spread of the MukB near the origin of replication along the two arms of the chromosome. Then we checked if the distribution over both arms was symmetric, by defining the local asymmetricity as the absolute difference between the fluorescent signal along the two chromosome halves divided by their sum (see Materials and methods). If the fluorescent intensity was equally spread over the two arms, the asymmetricity equals 0, while it equals 1 if all signal is on one of the chromosome arms (Fig. 2cd). In wildtype cells the median asymmetricity of the local DNA signal (which we used as the intrinsic measurement control) was 0.27 (Fig. 2ce), while for the MukB signal it was significantly higher at 0.33 (*N=284*) (Fig. 2de). Thus, it was clear that locally one of the arms (irrespective of the DNA content) within one chromosome had significantly more MukB signal compared to the other, although we could not identify which of the two arms is the left or right arm. Interestingly, when we measured the asymmetricity of the MukB in the ⊿*matP* cells, the spreading was even more asymmetric (median asymmetricity for DNA was 0.32, but for MukB 0.49, *N=108*) (Fig.2e). However, in the MukB^EQ^EF cells the asymmetricity of the DNA and MukB spreading along the chromosome was not significantly different (0.25 and 0.26, respectively *N=124)*.

**Figure 2.**
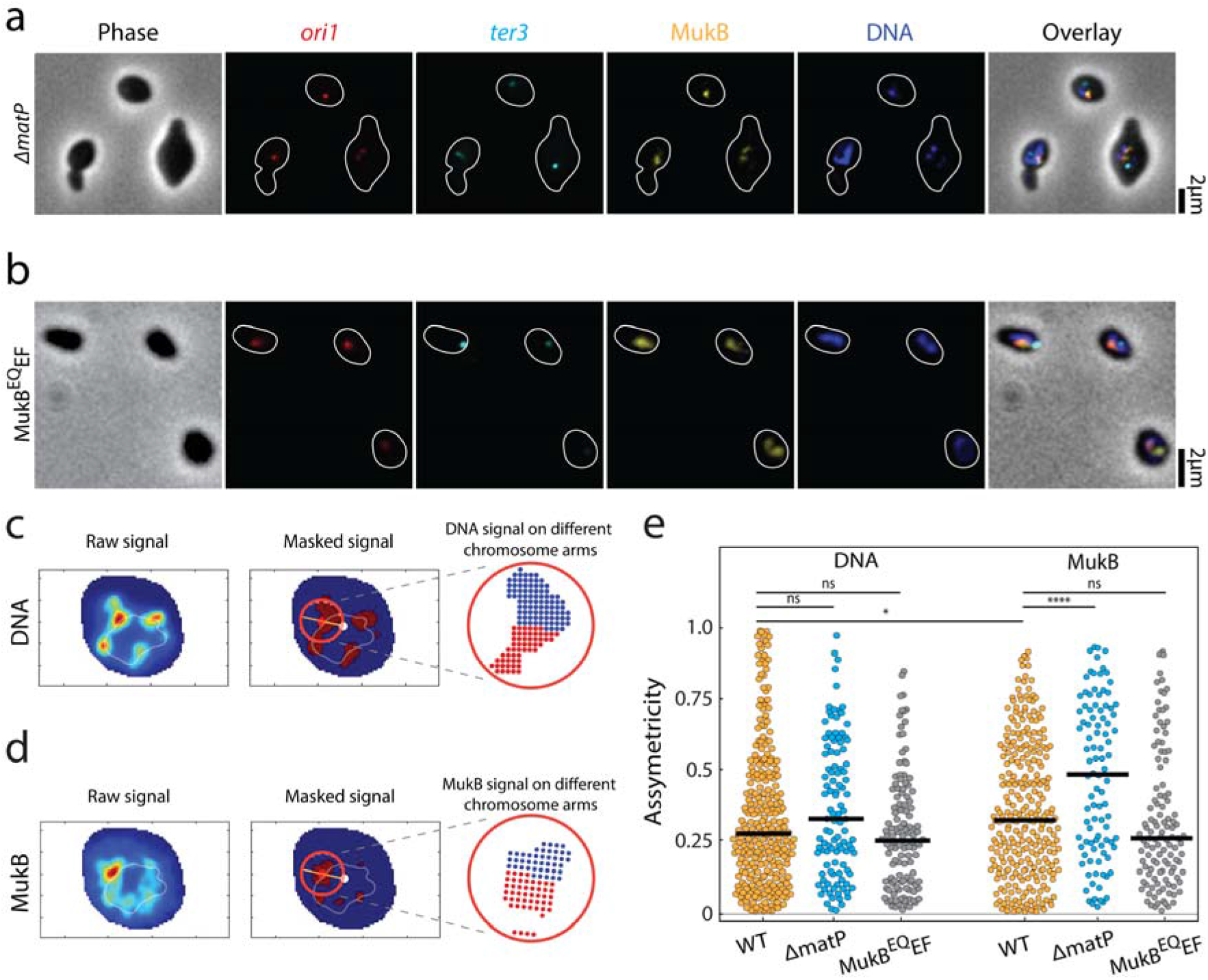
Asymmetric spreading of MukB complexes along the right and left arms of the chromosome. Representative images of **a**. ⊿*matP* and **b**. MukB^EQ^EF *E. coli* cells in phase contrast and the fluorescence channels of *ori1* (red), *ter3* (cyan), MukB (yellow), DNA stained with DAPI (blue) and their overlay. Cell outline is indicated in white. **c**. measuring the local spreading of DNA and **d**. MukB near the origin of replication. Left: the central ridge of the fluorescent signal is determined. Middle: the central axis (orange line) is detected along the origin (red spot) to the chromosome center of mass (white spot). Only for the circular area (red circle) around the origin, asymmetricity is determined. Right: a symmetry value is defined as the difference in signal from anticlockwise area (blue) and clockwise area (red) (relative to the line connecting ori and the center of mass) divided by their sum signal. Orange and light blue colors denote high and low signal intensity respectively. **e**. Asymmetricity distributions for DNA and MukB channels as defined in panels c and d for wildtype (*N= 284 cells*), ⊿*matP* (*N=108 cells*), and MukB^EQ^EF mutants (*N=124 cells*). Horizontal black lines represent median values. Statistical significance was determined by performing a single factor ANOVA test. The following conventions are used: ns: 0.05 < p, *: 0.01 < p < 0.05, ****: p < 0.0001. We report a significant difference in results if p < 0.05.

### MatP and ATP hydrolysis by MukBEF are needed for chromosome compaction and organization

To quantify chromosome compaction by MukBEF, we analyzed whether the DNA regions overlapping with MukBEF were associated with an increased DNA compaction compared to the rest of the chromosome. Interestingly, in wildtype cells the DNA density within the MukBEF regions was indeed higher, by a factor of 1.8 ± 0.05 (mean ± sem, *N=260 cells*), compared to the DNA signal elsewhere along the chromosome (Fig. 3a), highlighting MukBEF’s ability to locally compact the nucleoid. Despite the increased prevalence of MukBEF at the ter region in ⊿*matP* cells, the magnitude of local DNA compaction of MukBEF-occupied regions was not affected by the deletion of *matP*, as it was very similar to the wildtype (Fig. 3a; 1.75 ± 0.05 times higher (mean ± sem, *N=121 cells*)). Strikingly, the compaction of the DNA that overlapped with MukB in MukB^EQ^EF mutants was not significantly increased and only slightly higher compared to any other chromosomal region where MukBEF was not present (Fig. 3a; 1.2 ± 0.03 times higher than elsewhere along the chromosome (mean ± sem, *N=131 cells*)). This suggests that ATP hydrolysis is a requirement for compaction by MukBEF.

**Figure 3.**
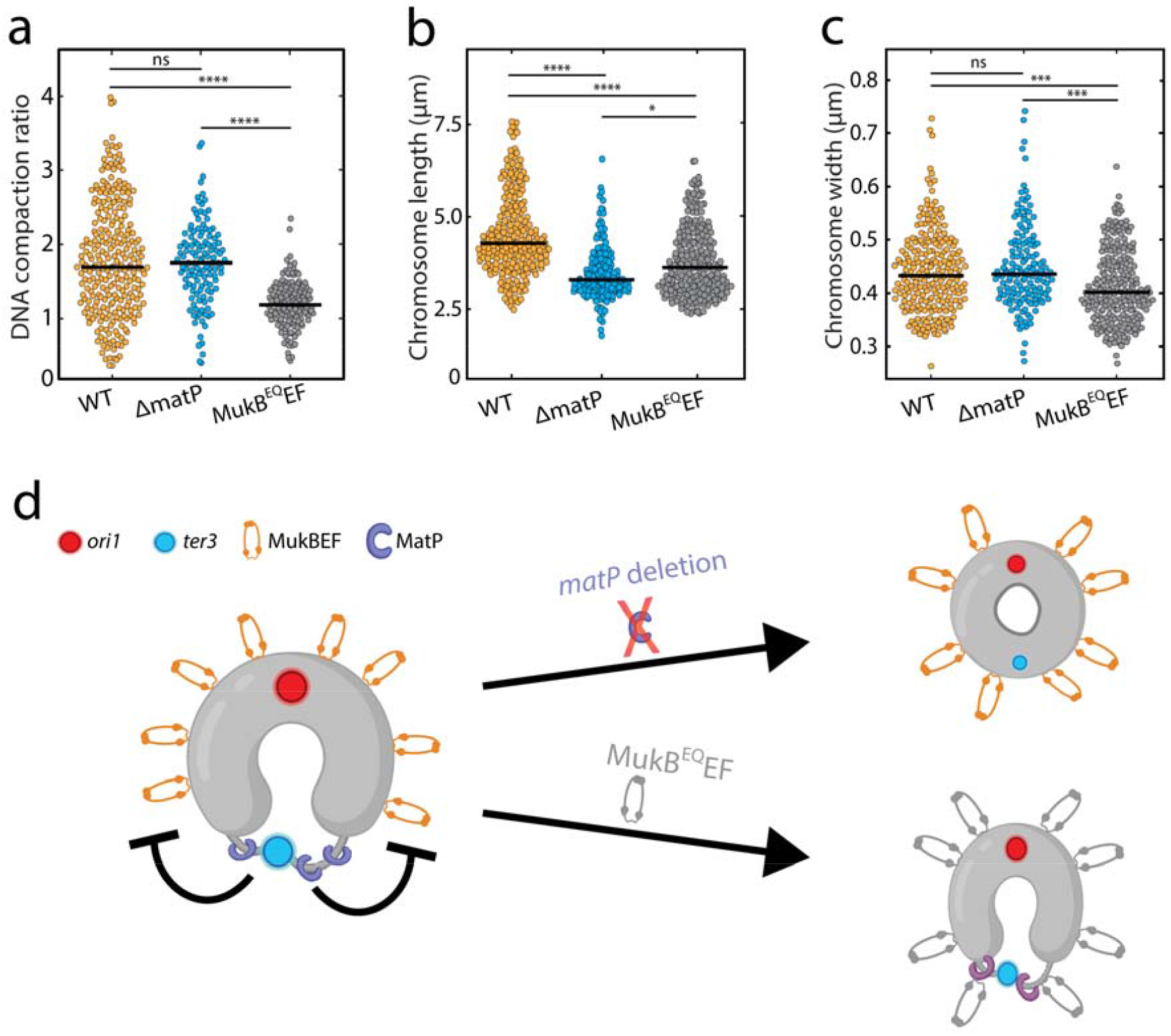
The effect of MukB on chromosome shape and compaction. **a**. Ratio of DNA density in the regions where MukB localizes versus elsewhere along the chromosome for wildtype (*N=260 cells*), ⊿*matP* (*N=121 cells*), and MukB^EQ^EF mutants (*N=131 cells*). **b**. Chromosome length distribution for wildtype (*N=222 cells*), ⊿*matP* (*N=153 cells*) and MukB^EQ^EF mutants (*N=226 cells*). **c**. Chromosome width distribution for wildtype (*N=222 cells*), ⊿*matP* (*N=153 cells*) and MukB^EQ^EF cells (*N=226 cells)*. **d**. Schematics depicting the circular nucleoid of *E. coli* with the position where MatP (purple) binds and a flexible decondensed terminus region (thin gray line). MukBEF is positioned away from the terminus near the origin of replication. Statistical significance was determined by performing a single factor ANOVA test. The following conventions are used: ns: 0.05 < p, *: 0.01 < p < 0.05, ***: 0.0001 < p < 0.001, ****: p < 0.0001. We report a significant difference in results if p < 0.05.

We next investigated the effects of altered MukBEF localization and activity on chromosome shape parameters. In both ⊿*matP* cells and MukB^EQ^EF mutants, we found that the chromosome contour length was significantly shorter than in wildtype (Fig. 3b). While wildtype cells had an average length of 4.5 ± 0.1 µm (mean ± sem, *N=222 cells*), it was reduced to 3.5 ± 0.1 µm (mean ± sem, *N=148 cells*) and 3.8 ± 0.1 µm (mean ±sem, *N=226 cells*) in ⊿*matP* cells and MukB^EQ^EF mutants, respectively. For the chromosome width, characterized by the average full-width-at-half-maximum along the chromosome (FWHM), a different trend was observed (Fig. 3c). We found that chromosomes of wildtype and ⊿*matP* cells had similar width of 0.44 ± 0.08 µm (mean ± sem, *N=222 cells*) and 0.45 ± 0.08 µm (mean ± sem, *N=148 cells*) respectively, while MukB^EQ^EF mutants formed a chromosome with a slightly reduced width of 0.41 ± 0.07 µm (mean ± sem, *N=226 cells*).

Since the DNA compaction in wildtype and ⊿*matP* mutant cells was similar, we conclude that MukBEF can bind and compact the chromosome independently of MatP protein. Aside from impeded MukBEF displacement over the DNA, the MukB^EQ^EF mutant also showed a clear impairment in its ability to compact the chromosome (Fig. 3a). Hence our findings suggest that MukBEF needs to hydrolyze ATP in order to compact the DNA and redistribute its position along the nucleoid (Fig. 3d), as similarly shown by others^13,35,36^. The altered chromosome length and width in the mutants show that MukBEF is an important factor in the global chromosome organization where both ATP hydrolysis and MukBEF’s interaction with MatP are required for its faithful functioning.

### *mukB* or *matP* deletion leads to chromosome decatenation defects and dimerization

Finally, to probe the roles of MukBEF and MatP in *E. coli* chromosome segregation, we visualized the chromosome structure after replication initiation in strains with either a *mukB* or *matP* deletion. However, since the *mukB* deleted cells did not grow at temperatures above 24°C^7^, cells could not be synchronized for replication initiation. As a result, these cells typically contained more than one chromosome at any given time. In 74% of the ⊿*mukB* cells (*N=172 cells*) there were 2 or more complete chromosomes, as recognized by a multitude of ori and ter foci, while this phenotype was not observed in wildtype cells^37^. As shown in Fig. 4a, chromosome dimers that were shaped like a figure-eight shape were observed in 91% of cells (*N= 172 cells*) with two complete chromosomes (i.e., cells with 2 ori and 2 ter foci), indicating that the decatenation of sister chromosomes was impaired in the absence of MukB. Interestingly, in the remaining 9% of polyploid cells, the chromosome dimers were organized in a toroidal configuration (Fig. 4b, more examples in Fig. S5).

**Figure 4.**
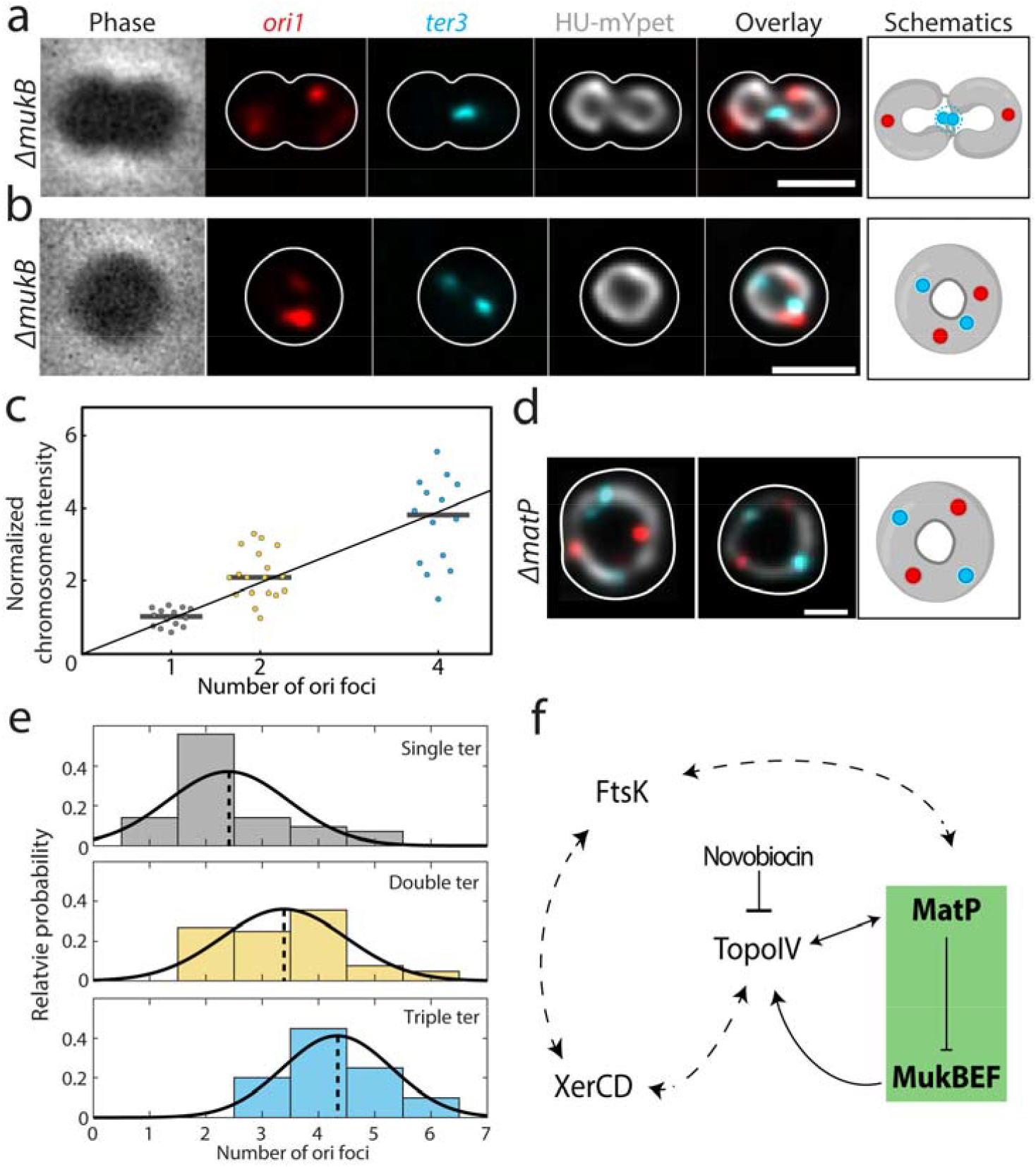
Impaired chromosome decatenation in ⊿*mukB* and ⊿*matP* cells. **a**. Phase contrast and fluorescence signals of *ori1* (RFP), *ter3* (CFP), DNA (HU-mYpet) and an overlay for a typical ⊿*mukB* cell widened with A22 are shown. The cell outline is shown in white and the impaired decatenation is schematically depicted. **b**. Idem as a, now for the donut-shaped chromosomes. **c**. Normalized sum chromosome signal (integrated DAPI signal) in ⊿*mukB* cells. Horizontal black lines show the median values (1,02; 2,10; 3,82) and a linear fit through these points and the origin is given (y=0.98x; R^2^=0.999). **d**. Distribution of the observed ori (x-axis) & ter (color) counts in ⊿*mukB* mutants (*N=167 cells*). Although observed, cells with four ter foci were excluded in the plot due to their low statistics (*N=5 cells*). The distributions are fitted with a Gausian function, whose mean is indicated by the dotted lines. **e**. Overlay of the fluorescence signals for DNA, ori and ter for two representative ⊿*matP* cells. The cell outline is shown in white, and the toroidal configuration is schematically depicted. **f**. Proposed model for the interaction between chromosomal organizers. Our data (green box) show that MatP inhibits chromosome compaction within the ter region by MukBEF, which interacts with TopoIV to fulfill its function. The interplay between TopoIV orchestrates chromosome decatenation, while novobiocin inhibits the latter, leading to segregation defects. These notions fit within the larger picture in which XerCD and FtsK are also interacting with TopoIV and MatP, respectively, as well as with each other. Double headed arrows indicate mutual regulation, while the dotted arrows indicate putative regulatory levels.

We related the number of ori foci to the total fluorescence intensity of the corresponding chromosome (normalized to cells with a single ori) and found that they scaled almost linearly (Fig. 4c), suggesting that the number of ori sites indeed indicated the number of fully replicated chromosomes in a cell. Dimer chromosomes had a normalized fluorescence intensity of 2.2 ± 0.7 (mean ± sd, *N=18* cells), while chromosomes with four oris had an intensity of 3.7 ± 1.2 (mean ± sd, *N=14* cells), compared to single chromosomes (1.0 ± 0.2 (mean ± sd, *N=14* cells)).

A closer inspection of the ori and ter foci in ⊿*mukB* cells revealed a striking difference compared to wildtype. Wildtype cells grown in minimal medium typically exhibited 2 ori/ter ratios: either we observed a single ori and a single ter (1:1) in cells with a single non-replicating chromosome, or we observed 2 oris and a single ter (i.e., a ratio of 2:1) in cells that were replicating. In ⊿*mukB* cells, however, only 17% of the cells showed either of these counts, whereas we found 15 other combinations of ori and ter counts (Fig. S5). We grouped these cells by the number of ter foci and plotted the ori counts in a histogram and found an increase in the average number of oris as the number of ters increases; from 2.4 ± 1.1 oris for cells with 1 ter, to 3.4 ± 1.1 for 2 ters and 4.3 ± 1.0 for 3 ters (all mean ± sd; Fig. 4d). The observation of single cells carrying four (or more) oris indicated that the next round of replication was initiated before sister chromosome decatenation^33^.

Deletion of *matP* also causes a strong phenotypic change in chromosome segregation. In ⊿*matP* cells, the ter region was not properly positioned relative to the mid cell throughout the segregation, as seen in wildtype. Cells with 2 ter foci were often observed in a stage of early septation, even though the circular sister chromosomes were still topologically catenated (Movie 1). Surprisingly, we could see the formation of toroidal dimer chromosomes in these cells as well. In 10.5% of the replicating ⊿*matP* cells, the two sister chromosomes would form one larger ring-like chromosome. Two pairs of ori and ter foci were present in these dimer chromosomes, which were evenly spaced and with a distinct order of ori-ter-ori-ter along the ring contour of the chromosome (Fig. 4e, more examples in Fig. S5). These cells displayed a similar phenotype as wildtype cells treated with the DNA gyrase inhibitor Novobiocin^38^ (Fig. S6), indicating that deletion of either *mukB* or *matP* impairs chromosome decatenation by Topo IV. Furthermore, both mutants showed impaired chromosome recombination and chromosome dimerization.

## Discussion

In this study, we investigated the role of MukBEF and MatP proteins in *E. coli* chromosome organization and segregation in A22-widened temperature-sensitive (*dnaCts*) *E. coli* cells. Because of the cell widening (typically twice larger in width and length compared to rod-shaped cells), the toroid-shape chromosome could be resolved and the positioning of the MukBEF along the nucleoid could be measured with increased spatial resolution without the need to overexpress MukBEF or cell fixation. Additionally, chromosome replication can be synchronized by first culturing the cells above the permissive temperatures (40°C), and subsequently re-initiating replication by transferring them to room temperature^28,29^.

Through quantitative imaging, we corroborated the earlier findings in rod-shaped cells that the MukBEF SMC in *E. coli* localizes near the origins of replication and away from the ter region^13^. In the absence of either ATP-hydrolysis by MukB (MukB^EQ^ mutant) or the MatP protein, we showed that MukBEF has a 3 to 5-fold higher association with ter, indicating that MukBEF normally binds to this macrodomain prior to being actively expelled toward ori by MatP (Fig. 1eg). Combined, the data elucidate that MatP-mediated MukBEF expulsion needs MukBEF’s ATP-dependent dissociation from the chromosome rather than preventing DNA binding altogether. This finding is in good agreement with earlier works, which showed that MukBEF is displaced from the ter region by MatP^15^.

Further, we observed an asymmetry of MukBEF occupancy on the left versus right arm of the chromosome (Fig. 2de), which is consistent with similar reports using HiC^39^, in other organisms such as *Corynebacterium glutamicum*^40^ and *Bacillus subtilis*^41^. There, however, SMCs were loading at specific *parS* genomic sequences near the origin of replication and spread preferentially towards the left arm of the chromosome, away from the highly transcribed ribosomal promoters positioned on the right arm. Our data are also consistent with Chip-seq on *E. coli* data showing a slight asymmetricity in the distribution of MukB binding along the chromosome arms, with higher preference for the right arm, once again away from the highly transcribed promoters^42^. The observation that the asymmetric spreading is lost in the MukB^EQ^EF cells points to the ATP-dependence of MukB spreading along the chromosome. We speculate that transcription-induced supercoiling provides physical barriers for the SMCs - impairing translocation over DNA or unbinding of the SMC -, resulting in an asymmetric spreading because highly transcribed promoters are unevenly distributed across the two arms.

Next, we showed that the local chromosomal regions that co-localize with MukBEF are on average almost twice as compacted as the rest of the chromosome (Fig. 3a). Again, MukBEF required its ATPase activity to carry out its DNA-compacting function, as the MukB^EQ^EF mutant showed no such increased compaction in MukBEF-occupied regions and had an altered chromosome width and length compared to wildtype (Fig. 3bc). This is consistent with the hypothesis that the local compaction is due to loop extrusion by MukBEF.

MukBEF and MatP are not merely factors that set the density of the chromosome, but they also play a major role in DNA segregation. For instance, MukBEF was recently shown to direct newly replicated origins towards the daughter cells^12^, and as a result, deletion of MukB resulted in anucleated cells and in cells that have unsegregated *oriC* at the older cell pole. MatP was conversely shown to be necessary for the proper localization of the ter domain after segregation and subsequent cell division^43^ and MatP was found to be involved in sister cohesion during the chromosomal movement right before cell division^44^. Here we demonstrated that segregation defects result from the disruption of the MatP or MukBEF function, after which *E. coli* chromosomes does adopt a toroidal dimer configuration (Fig. 4abe). In some cases, toroidal chromosomes consisting of two or more chromosomes formed, recognized by the multiple ori and ter foci (Fig. S5) and a higher total DNA sum signal (Fig. 4c). The variability in relative positioning of ori and ter foci along the chromosome in these cells might be the result of a disruption of recombination, where sister chromosomes remain concatenated and can move relative to each other. In the case of ⊿*mukB*, the fixed ori-ter-ori-ter order along the chromosome indicates the conjoinment of sisters into a single chromosome - probably through improper recombination between sister *dif* sites. These data are consistent with recent HiC data showing that both deletion of *mukB* or *matP*^45^, as well as extended sister-chromosome catenation through impairment of topo activity^46^ can lead to massive reorganization of the *E. coli* genome and to emergence of novel chromosome contact points.

As MukBEF and MatP themselves are not known to directly regulate sister chromosome decatenation, questions arise about other molecular players involved in the pathway to the observed segregation defects. The existing literature allows identification of potential interaction partners that may explain our observations (Fig. 4f). For example, loss of the ability to fully segregate chromosomes in ⊿*mukB* can at least in part be understood through the inherent loss of MukBEF’s ability to recruit chromosomal TopoIV, leading to reduced sister untangling throughout segregation^22^. Further, MatP was found to be essential for DNA translocation by the FtsK protein^47^, which is another component that processes DNA in the late stages of cell division^48^. The FtsK-mediated translocation at *dif* ultimately ceases upon contact with the XerCD recombinase system^49,50^, and XerCD and TopoIV decatenate the sister chromosomes in the final phase of segregation^51–53^. So, there are multiple observations that exemplify MatP’s involvement in the tight regulation of TopoIV-mediated sister-resolution at ter: MatP’s competition with TopoIV to bind MukB^19^, its function to actively displace MukBEF from ter, and the XerCD-TopoIV interaction. XerCD necessitates and specifically guides TopoIV to *dif* to resolve concatenated sisters^54^. *matP* deletion can therefore disrupt chromosome recombination, leading to a single toroidal dimer chromosome instead of properly decatenated sisters. In the absence of MatP, the MukBEF-TopoIV complex spreads to the ter region, where XerCD normally orchestrates sister decatenation by guiding TopoIV to a specific site. The increased abundance of MukBEF-TopoIV in this region might interfere with this site-specific XerCD action, thereby inducing uncontrolled sister decatenation and concatenation by TopoIV with the observed population of conjoined dimers as a result of this dysregulated recombination. Predominant MatP binding to the hinge of TopoIV in the *matS*-MatP-enriched ter could minimize random action of TopoIV in wildtype cells.

All in all, applicability of our approach to resolve protein-chromosome spatial interactions in widened live *E. coli* cells showed new insights in how MukBEF and MatP shape the chromosome. By performing quantitative fluorescence microscopy on artificially enlarged cells, we effectively gained spatial resolution of the interactions between multiple simultaneously tagged targets. This approach allowed us to acquire new information on how MukBEF and MatP localize relative to each other and along the genome. The presented platform holds good potential to further resolve the spatial interactome that governs chromosome homeostasis, replication, and segregation in prokaryotes.

## Materials and methods

### Strain construction

All strains were derivatives of *E. coli* K12 AB1157 strain and were constructed by P1 transduction^55^. To construct strain AJ2820, strain SN192^13^ (*lacO240::hyg at ori1, tetO240::gen at ter3, Plac-lacImCherry frt at leuB, Plac-tetR-mCerulean frt at galK, mukB-mYPet frt*), was transduced with P1 phage derived from FW1957^17^ (*dnaC2(ts)* ⊿*mdoB::aph :: frt*) to result in a DnaC temperature sensitivity.

To construct strain AJ2822(⊿*matP*), strain SN302^13^ (*lacO240::hyg at ori1, tetO240::gen at ter3, Plac-lacImCherry frt at leuB, Plac-tetR-mCerulean frt at galK, mukB-mYPet frt*, ⊿*matP::cat CM*^*R*^), a kind gift from David Sherratt, was transduced with P1 phage derived from FW1957^17^ (*dnaC2(ts)* ⊿*mdoB::aph :: frt*) to result in a DnaC temperature sensitivity. ⊿*mukB* (strain Ab243) and MukB^EQ^EF (strain SN311) were generated elsewhere^13^.

### Growth conditions

For obtaining cells with circular chromosomes, we grew cells in liquid M9 minimal medium (Fluka Analytical) supplemented with 2 mM MgSO_4_, 0.1mM CaCl_2_, 0.4% glycerol (Sigma-Aldrich), and 0.01% PHA (Fluka Analytical) overnight at 30°C to reach late exponential phase. On the day of the experiment, the overnight culture was refreshed (1:100 vol) for 2 hours on fresh M9 minimal medium at 30 °C. We then pipetted 1μl culture onto a cover glass and immediately covered the cells with a flat agarose pad, containing the above composition of M9 medium, A22 (final 4μg/ml), as well as 3% agarose. The cover glass was then placed onto a baseplate and sealed with parafilm to prevent evaporation. The baseplate was placed onto the microscope inside a heated chamber set at 40°C for 2.5 hours to stop the cells from replicating and to let them grow into round shapes. In ⊿*mukB* cells incubation and imaging were performed at room temperature (22 °C) as these cells are unable to grow at higher temperatures.

For experiments with cells lacking HU-mYPet labeling (AB243, AJ2820 & AJ2822) we used DAPI to stain the nucleoids. On the day of the experiment, the overnight culture was refreshed (1:100 vol) for 2 hours on fresh M9 minimal medium at 30°C. We then added A22 (final 4μg/ml) to the cell culture and transferred the sample to a 40°C incubator for 2.5hours. To minimize any possible artifacts arising from the incubation of cells with the DAPI^56^ (Fig. S7), we added DAPI (final concentration 1µg/ml) last, just before imaging. 1ml of the cell sample was incubated with DAPI for less than 1 min and then centrifuged at 4000 rpm for 2 minutes. 900μl of the supernatant was removed and the pellet was resuspended in the remaining ∼100μl medium. 1μl of the remaining culture was deposited onto the cover glass immediately covered with a flat agarose pad, containing M9 medium, A22 (final 4μg/ml), as well as 3% agarose. The sample was placed onto the microscope-stage and imaged immediately.

For treatment of replicating cells with Novobiocin^38^, we first grew the cells in the presence of A22 as described above for 2.5 h to ensure they reach desired size and shape. Then we moved the baseplate to room-temperature for 10 min to re-initiate replication and afterwards added 10 μl of Novobiocin (∼50 μg/ml final) to the agarose pad during replication initiation phase. Finally, the cells were moved back to 40 °C chamber and imaged.

### Fluorescence imaging

Wide-field Z scans were carried out using a Nikon Ti-E microscope with a 100X CFI Plan Apo Lambda Oil objective with an NA of 1.45. The field of view corresponded to 2048 × 2048 pixels with a pixel size of 0.065 μm x 0.065 μm. The microscope was enclosed by a custom-made chamber that was pre-heated overnight and kept at 40°C (except when imaging the ⊿*mukB* cells). mCerulean was excited by SpectraX LED (Lumencor) λ_ex_ = 430-450 through a CFP filter cube (λ_ex_ / λ_bs_ / λ_em_ =426-446 / 455 / 460-500 nm). mYPet signal was excited by SpectraX LED λ_ex_ = 510/25 nm through a triple band-pass filter λ_em_ = 465/25 – 545/30 – 630/60 nm. mCherry signal was excited by SpectraX LED λ_ex_ = 575/25 through the same triple band-pass filter. Fluorescent signals were captured by Andor Zyla USB3.0 CMOS Camera. For each channel, between 3-19 slices were taken with a vertical step size of 227 nm (up to 2.3 μm in total).

### Image deconvolution

Image stacks of 3-19 slices of Z stack in wide-field imaging were deconvolved using the Huygens Professional deconvolution software (Scientific Volume Imaging, Hilversum, The Netherlands), using an iterative Classic Maximum Likelihood Estimate (CMLE) algorithm with a point spread function (PSF) experimentally measured using 200 nm multicolor Tetrabeads (Invitrogen). The PSF of the single-frame non-deconvolved widefield images had a FWHM of 350 nm horizontally and 800 nm vertically. Deconvolution reduced the out-of-focus noise in the images, which also led to an improvement in lateral resolution.

### Automated cell identification

Phase contrast images were fed into a customized Matlab program described earlier^28^, to produce masks of cell boundaries, which then were used to allocate chromosomes and foci in other fluorescence channels. A manual correction and rejection process was carried out as a final step of quality control, to correct or reject cells when neighboring cells were too close to allow the automated program to be distinguished.

### Compaction, colocalization, foci counting and asymmetricity

First, a customized Matlab program was used to threshold all fluorescence signals with a Gaussian filter and remove background signal. The fluorescence intensity of the DNA signal was used as a direct measure for its density and relative compaction. We defined MukB-DNA colocalization as the percentage of the total DNA intensity that overlaps with the MukB mask. As both ori and ter are single spots in approximation, we made colocalization of MukB with the foci binary. Either (part of) a locus overlapped with MukB, which we classified as colocalization or there was no overlap and thus no colocalization. The ori and ter foci were counted for each cell and cells with more ter than ori foci were discarded.

To measure the asymmetricity of DNA and MukBEF signal, first the fluorescence intensities and the central ridge of the chromosome toroid were measured. Next, the orientation was set by measuring a central axis through the position of the origin of replication and the center-of-mass of the chromosome. Only a circular area around the origin was evaluated for local symmetry. The diameter of the masking circle was 20 pixels = 1.3 µm, in order to encompass the typical chromosome width and the typical local chromosome cluster sizes (Fig. S4). The local asymmetricity *A* for the DNA and MukB channels was defined by the following formula:

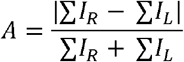

Where ∑*I*_*R*_ and ∑*I*_*L*_ are the sum fluorescent signals (DNA and MukB) on the right and left sides of the chromosome respectively. If the fluorescent intensity is equally spread, the asymmetricity is equal to 0, if all signal is on one of the chromosome arms, then it is equal to1. Due to the unknown orientation of the cell, the colors (red and blue on Figure 2d) cannot be attributed to the genomic orientation of left and right chromosome arms.

## Data availability

The data underlying Figure 1e, 1f, 3a,3b,3c, 4c and 4e are provided as a Source Data file. Datasets that were acquired and analyzed during the current study are available from the corresponding author upon request.

## Code availability

The analysis codes that were used in the current study are available from the corresponding author upon request.

## Acknowledgments

Financial support was provided by the ERC Advanced Grant no. 883684 and the NanoFront and BaSyC programs of NWO/OCW, and by the Swiss National Science Foundation (P300P2_177768). We thank David Sherratt for the kind gift of strains, Miloš Tišma and Huyen My Nguyen for discussions. Schematics on Fig.1g, Fig.3d & Fig.4a, b, e were created with BioRender.com.

## Notes

### Competing Interest Statement

The authors have declared no competing interest.

## Bibliography

1. Travers, A. & Muskhelishvili, G. Bacterial chromatin. Curr. Opin. Genet. Dev. 15, 507–514 (2005) DOI:10.1016/J.GDE.2005.08.006.

2. Blot, N., Mavathur, R., Geertz, M., Travers, A. & Muskhelishvili, G. Homeostatic regulation of supercoiling sensitivity coordinates transcription of the bacterial genome. EMBO Rep. 7, 710–715 (2006) DOI:10.1038/SJ.EMBOR.7400729.

3. Luijsterburg, M. S., Noom, M. C., Wuite, G. J. L. & Dame, R. T. The architectural role of nucleoid-associated proteins in the organization of bacterial chromatin: A molecular perspective. J. Struct. Biol. 156, 262–272 (2006) DOI:10.1016/J.JSB.2006.05.006.

4. Reyes-Lamothe, R., Wang, X. & Sherratt, D. Escherichia coli and its chromosome. Trends Microbiol. 16, 238–245 (2008) DOI:10.1016/j.tim.2008.02.003.

5. Thanbichler, M., Viollier, P. H. & Shapiro, L. The structure and function of the bacterial chromosome. Curr. Opin. Genet. Dev. 15, 153–162 (2005) DOI:10.1016/j.gde.2005.01.001.

6. Nolivos, S. & Sherratt, D. The bacterial chromosome: Architecture and action of bacterial SMC and SMC-like complexes. FEMS Microbiol. Rev. 38, 380–392 (2014) DOI:10.1111/1574-6976.12045.

7. Niki, H., Jaffé, A., Imamura, R., Ogura, T. & Hiraga, S. The new gene mukB codes for a 177 kd protein with coiled-coil domains involved in chromosome partitioning of E. coli. EMBO J. 10, 183–193 (1991).

8. Petrushenko, Z. M., Lai, C. H. & Rybenkov, V. V. Antagonistic interactions of kleisins and DNA with bacterial condensin MukB. J. Biol. Chem. (2006) DOI:10.1074/jbc.M606723200.

9. Rybenkov, V. V., Herrera, V., Petrushenko, Z. M. & Zhao, H. MukBEF, a chromosomal organizer. J. Mol. Microbiol. Biotechnol. 24, 371–383 (2014) DOI:10.1159/000369099.

10. Chen, N. et al. ATP-induced shrinkage of DNA with MukB protein and the MukBEF complex of escherichia coli. J. Bacteriol. 190, 3731–3737 (2008) DOI:10.1128/JB.01863-07.

11. Mäkelä, J. & Sherratt, D. J. Organization of the Escherichia coli Chromosome by a MukBEF Axial Core. Mol. Cell 78, 250–260 (2020) DOI:10.1016/J.MOLCEL.2020.02.003.

12. Mäkelä, J., Uphoff, S. & Sherratt, D. J. Nonrandom segregation of sister chromosomes by Escherichia coli MukBEF. Proc. Natl. Acad. Sci. U. S. A. 118, (2021) DOI:10.1073/PNAS.2022078118/SUPPL_FILE/PNAS.2022078118.SM03.MP4.

13. Nolivos, S. et al. MatP regulates the coordinated action of topoisomerase IV and MukBEF in chromosome segregation. Nat. Commun. 7, 10466 (2016) DOI:10.1038/ncomms10466.

14. Espéli, O. et al. A MatP–divisome interaction coordinates chromosome segregation with cell division in E. coli. EMBO J. 31, 3198–3211 (2012) DOI:10.1038/EMBOJ.2012.128.

15. Mercier, R. et al. The MatP/matS Site-Specific System Organizes the Terminus Region of the E. coli Chromosome into a Macrodomain. Cell 135, 475–485 (2008) DOI:10.1016/j.cell.2008.08.031.

16. Bürmann, F., Funke, L. F. H., Chin, J. W. & Löwe, J. Cryo-EM structure of MukBEF reveals DNA loop entrapment at chromosomal unloading sites. Mol. Cell 81, 4891-4906.e8 (2021) DOI:10.1016/j.molcel.2021.10.011.

17. Danilova, O., Reyes-Lamothe, R., Pinskaya, M., Sherratt, D. & Possoz, C. MukB colocalizes with the oriC region and is required for organization of the two Escherichia coli chromosome arms into separate cell halves. Mol. Microbiol. 65, 1485–1492 (2007) DOI:10.1111/j.1365-2958.2007.05881.x.

18. Yamazoe, M. et al. Complex formation of MukB, MukE and MukF proteins involved in chromosome partitioning in Escherichia coli. EMBO J. 18, 5873–5884 (1999) DOI:10.1093/emboj/18.21.5873.

19. Nicolas, E. et al. The SMC complex MukBEF recruits topoisomerase IV to the origin of replication region in live Escherichia coli. MBio 5, e01001–13 (2014) DOI:10.1128/mBio.01001-13.

20. Deweese, J. E. & Osheroff, N. The DNA cleavage reaction of topoisomerase II: Wolf in sheep’s clothing. Nucleic Acids Res. 37, 738–748 (2009) DOI:10.1093/nar/gkn937.

21. Ashley, R. E. et al. Activities of gyrase and topoisomerase IV on positively supercoiled DNA. Nucleic Acids Res. 45, 9611–9624 (2017) DOI:10.1093/nar/gkx649.

22. Fisher, G. L. et al. Competitive binding of MatP and topoisomerase IV to the MukB hinge domain. Elife 10, 1–25 (2021) DOI:10.7554/eLife.70444.

23. Goto, T. & Wang, J. C. Yeast DNA topoisomerase II. An ATP-dependent type II topoisomerase that catalyzes the catenation, decatenation, unknotting, and relaxation of double-stranded DNA rings. J. Biol. Chem. 257, 5866–5872 (1982).

24. Peng, H. & Marians, K. J. Decatenation activity of topoisomerase IV during oriC and pBR322 DNA replication in vitro. Proc. Natl. Acad. Sci. U. S. A. 90, 8571–8575 (1993) DOI:10.1073/pnas.90.18.8571.

25. Seol, Y., Hardin, A. H., Strub, M. P., Charvin, G. & Neuman, K. C. Comparison of DNA decatenation by Escherichia coli topoisomerase IV and topoisomerase III: Implications for non-equilibrium topology simplification. Nucleic Acids Res. 41, 4640–4649 (2013) DOI:10.1093/nar/gkt136.

26. Saifi, B. & Ferat, J. L. Replication fork reactivation in a dnaC2 mutant at non-permissive temperature in escherichia coli. PLoS One 7, e33613 (2012) DOI:10.1371/journal.pone.0033613.

27. Varma, A. & Young, K. D. In Escherichia coli, MreB and FtsZ direct the synthesis of lateral cell wall via independent pathways that require PBP 2. J. Bacteriol. 191, 3526–3533 (2009) DOI:10.1128/JB.01812-08.

28. Wu, F. et al. Direct imaging of the circular chromosome in a live bacterium. Nat. Commun. 10, 1–9 (2019) DOI:10.1038/s41467-019-10221-0.

29. Japaridze, A., Gogou, C., Kerssemakers, J. W. J., Nguyen, H. M. & Dekker, C. Direct observation of independently moving replisomes in Escherichia coli. Nat. Commun. 11, 1–10 (2020) DOI:10.1038/s41467-020-16946-7.

30. Karczmarek, A. et al. DNA and origin region segregation are not affected by the transition from rod to sphere after inhibition of Escherichia coli MreB by A22. Mol. Microbiol. 65, 51– 63 (2007) DOI:10.1111/J.1365-2958.2007.05777.X.

31. Mickiewicz, K. M. et al. Possible role of L-form switching in recurrent urinary tract infection. Nat. Commun. 10, 1–9 (2019) DOI:10.1038/s41467-019-12359-3.

32. Wang, X., Liu, X., Possoz, C. & Sherratt, D. J. The two Escherichia coli chromosome arms locate to separate cell halves. Genes Dev. 20, 1727–1731 (2006) DOI:10.1101/GAD.388406.

33. Khan, S. R., Mahaseth, T., Kouzminova, E. A., Cronan, G. E. & Kuzminov, A. Static and dynamic factors limit chromosomal replication complexity in Escherichia coli, avoiding dangers of runaway overreplication. Genetics 202, 945–960 (2016) DOI:10.1534/genetics.115.184697.

34. Wu, F. et al. Cell Boundary Confinement Sets the Size and Position of the E. coli Chromosome. Curr. Biol. 29, 2131–2144 (2019) DOI:10.1016/J.CUB.2019.05.015.

35. Woo, J. S. et al. Structural Studies of a Bacterial Condensin Complex Reveal ATP-Dependent Disruption of Intersubunit Interactions. Cell 136, 85–96 (2009) DOI:10.1016/J.CELL.2008.10.050.

36. Badrinarayanan, A., Reyes-Lamothe, R., Uphoff, S., Leake, M. C. & Sherratt, D. J. In vivo architecture and action of bacterial structural maintenance of chromosome proteins. Science 338, 528–531 (2012) DOI:10.1126/SCIENCE.1227126/SUPPL_FILE/BADRINARAYANAN.SM.PDF.

37. Nordman, J., Skovgaard, O. & Wright, A. A novel class of mutations that affect DNA replication in E. coli. Mol. Microbiol. 64, 125–138 (2007) DOI:10.1111/j.1365-2958.2007.05651.x.

38. Maxwell, A. The interaction between coumarin drugs and DNA gyrase. Mol. Microbiol. 9, 681–686 (1993) DOI:10.1111/j.1365-2958.1993.tb01728.x.

39. Dostie, J. et al. Chromosome Conformation Capture Carbon Copy (5C): A massively parallel solution for mapping interactions between genomic elements. Genome Res. 16, 1299–1309 (2006) DOI:10.1101/GR.5571506.

40. Böhm, K. et al. Chromosome organization by a conserved condensin-ParB system in the actinobacterium Corynebacterium glutamicum. Nat. Commun. 11, 1–17 (2020) DOI:10.1038/s41467-020-15238-4.

41. Antar, H. et al. Relief of ParB autoinhibition by parS DNA catalysis and recycling of ParB by CTP hydrolysis promote bacterial centromere assembly. Sci. Adv. 7, (2021) DOI:10.1126/SCIADV.ABJ2854/SUPPL_FILE/SCIADV.ABJ2854_SM.PDF.

42. Verma, S. C., Qian, Z. & Adhya, S. L. Architecture of the Escherichia coli nucleoid. PLOS Genet. 15, e1008456 (2019) DOI:10.1371/JOURNAL.PGEN.1008456.

43. Galli, E., Midonet, C., Paly, E. & Barre, F. X. Fast growth conditions uncouple the final stages of chromosome segregation and cell division in Escherichia coli. PLoS Genet. 13, e1006702 (2017) DOI:10.1371/JOURNAL.PGEN.1006702.

44. Crozat, E. et al. Post-replicative pairing of sister ter regions in Escherichia coli involves multiple activities of MatP. Nat. Commun. 11, 1–12 (2020) DOI:10.1038/s41467-020-17606-6.

45. Lioy, V. S. et al. Multiscale Structuring of the E. coli Chromosome by Nucleoid-Associated and Condensin Proteins. Cell 172, 771–783 (2018) DOI:10.1016/j.cell.2017.12.027.

46. Conin, B. et al. Extended sister-chromosome catenation leads to massive reorganization of the E. coli genome. Nucleic Acids Res. 50, 2635–2650 (2022) DOI:10.1093/NAR/GKAC105.

47. Stouf, M., Meile, J. C. & Cornet, F. FtsK actively segregates sister chromosomes in Escherichia coli. Proc. Natl. Acad. Sci. U. S. A. 110, 11157–11162 (2013) DOI:10.1073/PNAS.1304080110/-/DCSUPPLEMENTAL.

48. Wang, M., Fang, C., Ma, B., Luo, X. & Hou, Z. Regulation of cytokinesis: FtsZ and its accessory proteins. Curr. Genet. 66, 43–49 (2020) DOI:10.1007/s00294-019-01005-6.

49. Bonné, L., Bigot, S., Chevalier, F., Allemand, J. F. & Barre, F. X. Asymmetric DNA requirements in Xer recombination activation by FtsK. Nucleic Acids Res. 37, 2371–2380 (2009) DOI:10.1093/NAR/GKP104.

50. Graham, J. E., Sivanathan, V., Sherratt, D. J. & Arciszewska, L. K. FtsK translocation on DNA stops at XerCD-dif. Nucleic Acids Res. 38, 72–81 (2010) DOI:10.1093/NAR/GKP843.

51. El Sayyed, H. et al. Mapping Topoisomerase IV Binding and Activity Sites on the E. coli Genome. PLoS Genet. 12, 1–22 (2016) DOI:10.1371/journal.pgen.1006025.

52. Farrokhi, A., Liu, H. & Szatmari, G. Characterization of the Chromosome Dimer Resolution Site in Caulobacter crescentus. J. Bacteriol. 201, e00391–19 (2019) DOI:10.1128/JB.00391-19.

53. Sciochetti, S. A. & Piggot, P. J. A tale of two genomes: resolution of dimeric chromosomes in Escherichia coli and Bacillus subtilis. Res. Microbiol. 151, 503–511 (2000) DOI:10.1016/S0923-2508(00)00220-5.

54. Gogou, C., Japaridze, A. & Dekker, C. Mechanisms for Chromosome Segregation in Bacteria. Front. Microbiol. 12, 685687 (2021) DOI:10.3389/FMICB.2021.685687/BIBTEX.

55. Thomason, L. C., Costantino, N. & Court, D. L. E. coli Genome Manipulation by P1 Transduction. in Current Protocols in Molecular Biology (2014). DOI:10.1002/0471142727.mb0117s79.

56. Japaridze, A., Benke, A., Renevey, S., Benadiba, C. & Dietler, G. Influence of DNA binding dyes on bare DNA structure studied with atomic force microscopy. Macromolecules 48, 1860– 1865 (2015) DOI:10.1021/ma502537g.

