## Supplemental FIgures for "MukBEF-dependent chromosomal organization in widened *Escherichia coli*"

**SI Figures**

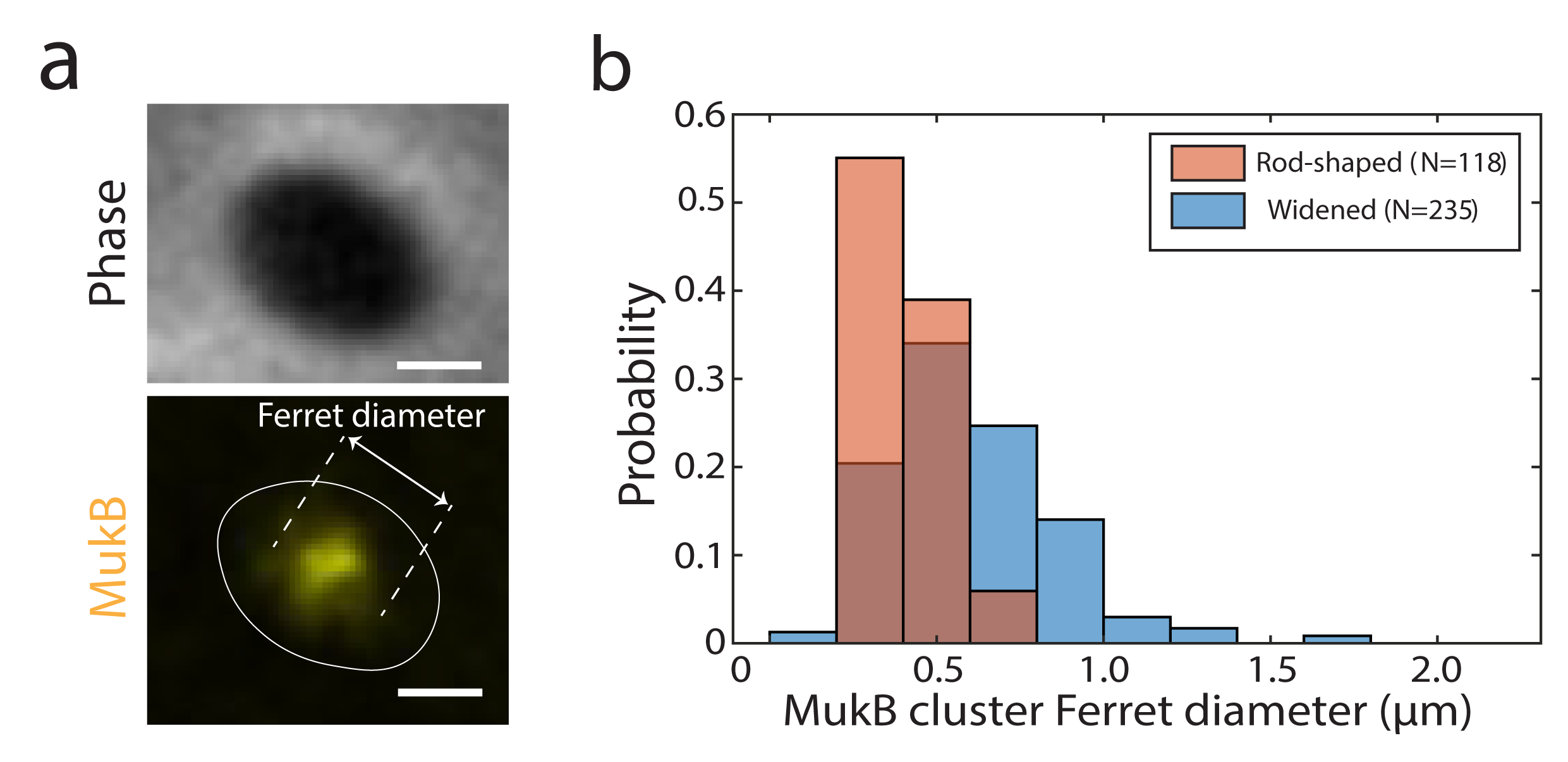

**Figure S1. a.** Representative image of a widened *E. coli* cell in phase contrast (top) and the MukB-YFP fluorescence channel (bottom). Cell outline is indicated in white. Scale bar is 1µm and the Ferret diameter of the MukB cluster is indicated. **b.** Ferret diameter distribution of MukB clusters in rod-shaped cells (red, *N=118*) and widened *E.coli* cells (blue, *N=235*). MukB clusters were significantly larger in widened cells compared to rod-shaped cells (p < 0.0001 with a single factor ANOVA test).

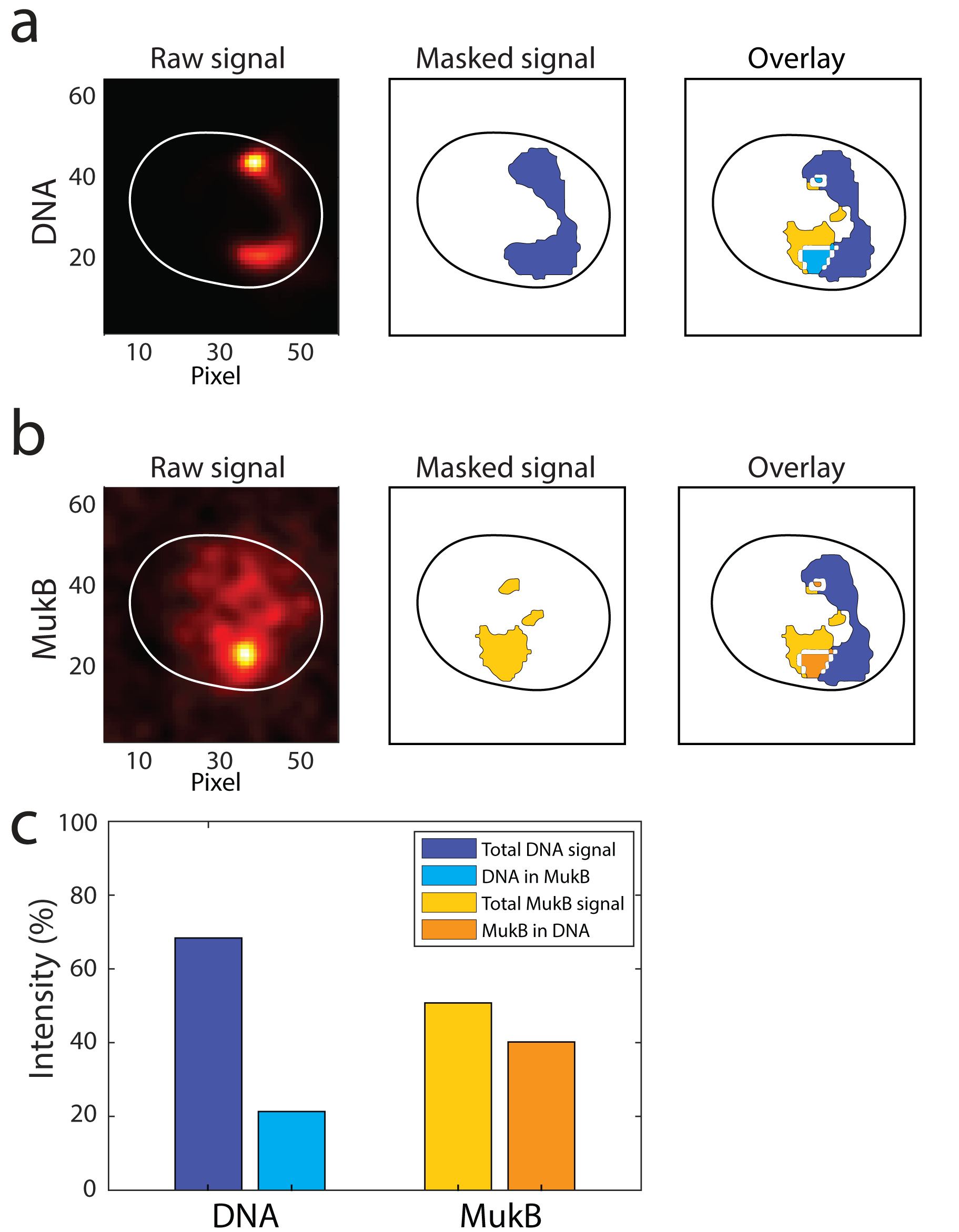

**Figure S2.** Quantitative co-localization of MukBEF signal with DNA signal. Fluorescent images of **a.** DNA and **b.** MukB channel. First the fluorescent signals are masked and then the masked signal of both channels is overlaid. Pixel size is 0.065 μm x 0.065 μm. Cell outline is indicated with white or black continuous lines. **c.** The relative intensity overlap between the two channels (DNA and MukB) is calculated, by determining the fraction of total fluorescence intensity of one channel that lies within the masked region of the other channel.

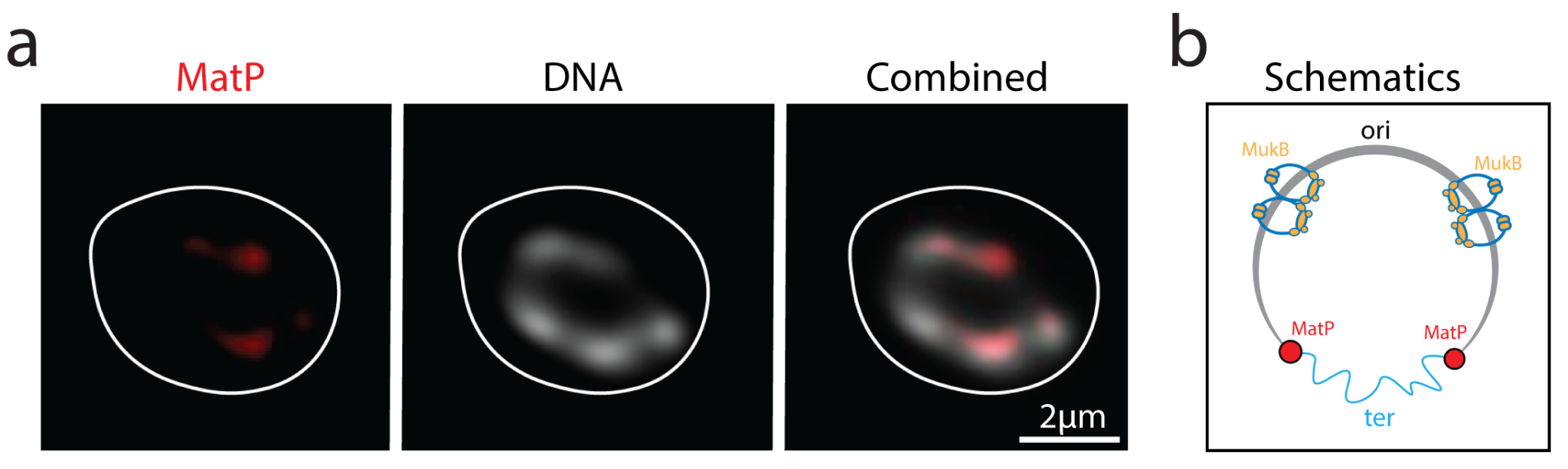

**Figure S3. a.** Localization of MatP in widened cells. MatP (tagged with mCherry) (left), DNA (HU-mYpet) (middle) and an overlay of the two channels (right). Cell outline is indicated in white. **b.** Schematics depicting the circular nucleoid of *E. coli* with the position where MatP (red dots) bind and the flexible decondensed terminus region (blue wiggly line). MukBEF (dark blue and yellow) is positioned away from the terminus near the origin or replication.

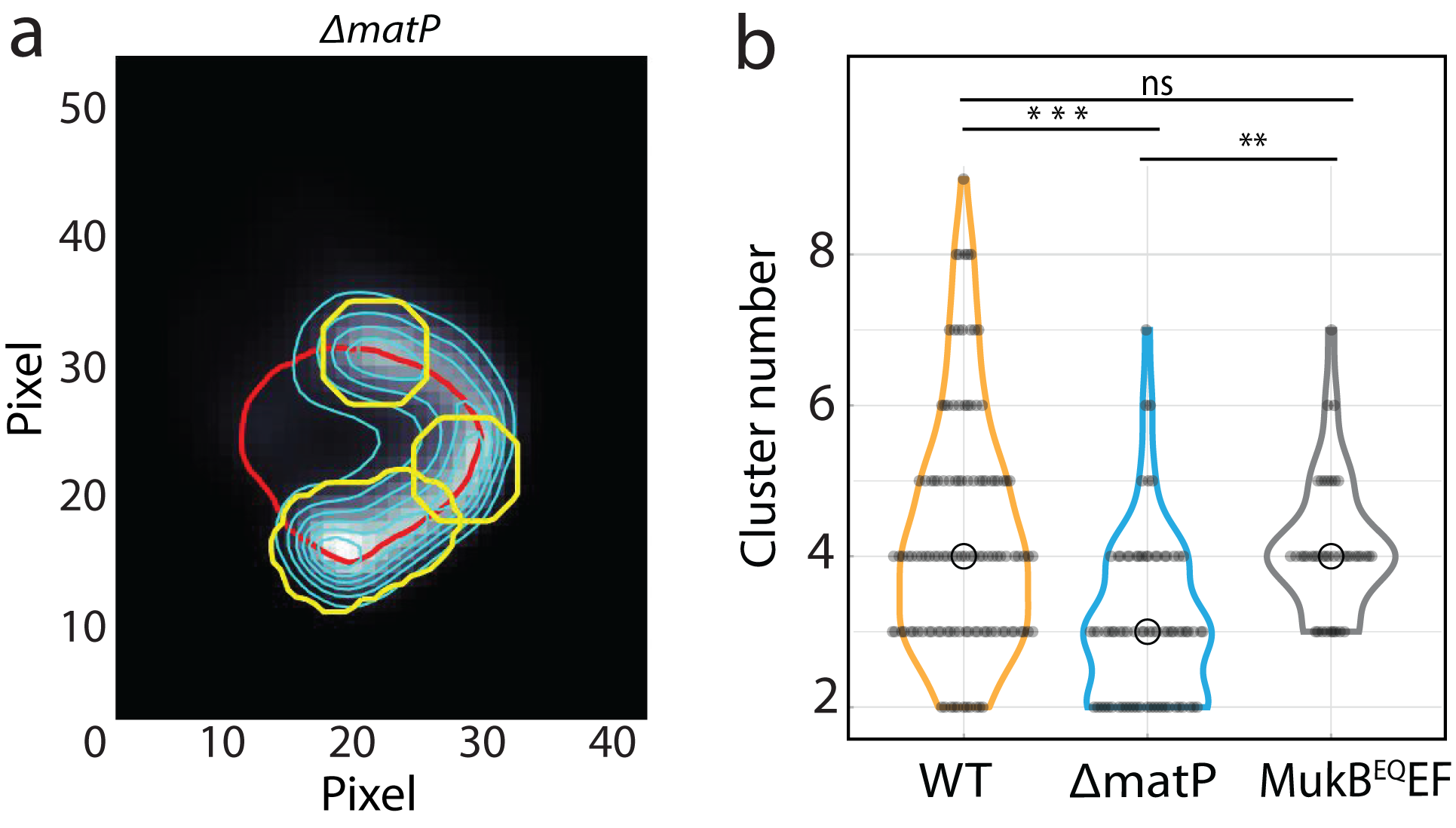

**Figure S4. a.** Typical image of a chromosome of a *ΔmatP* cell after performing cluster analysis. The chromosome central ridge is shown in red line and the blue lines represent the intensity contour lines. Yellow lines define DNA cluster contours. Pixel size is 0.065 μm x 0.065 μm. **b.** Violin plot of the number of DNA clusters for various cell lines (wildtype in yellow, *ΔmatP* in blue, MukB^EQ^EF in grey). Black circles show the median values. Statistical significance was determined by performing a single factor ANOVA test. The following conventions are used: ns: 0.05 < p, **: 0.001 < p < 0.01, ***: 0.0001 < p < 0.001.

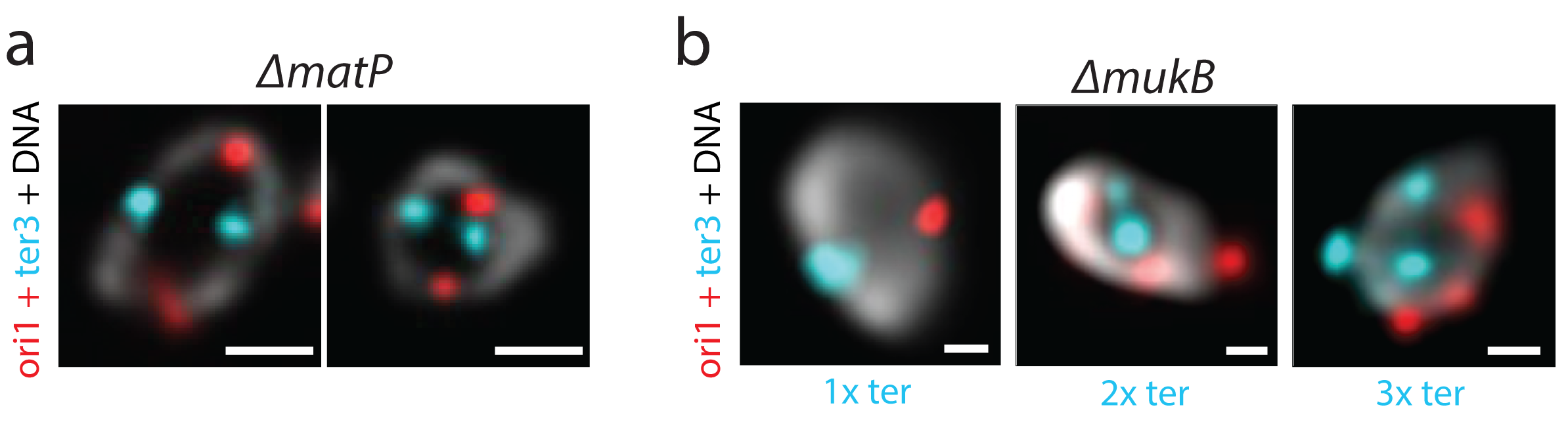

**Figure S5. a.** Typical microscopy images of dimer chromosomes (with two ori (red) and two ter (cyan) foci) in *ΔmatP* cells. **b.** Typical microscopy images of chromosomes in *ΔmukB* cells. Chromosomes display single, double or triple ori (red) and ter (cyan) foci. Scale bars are 1um.

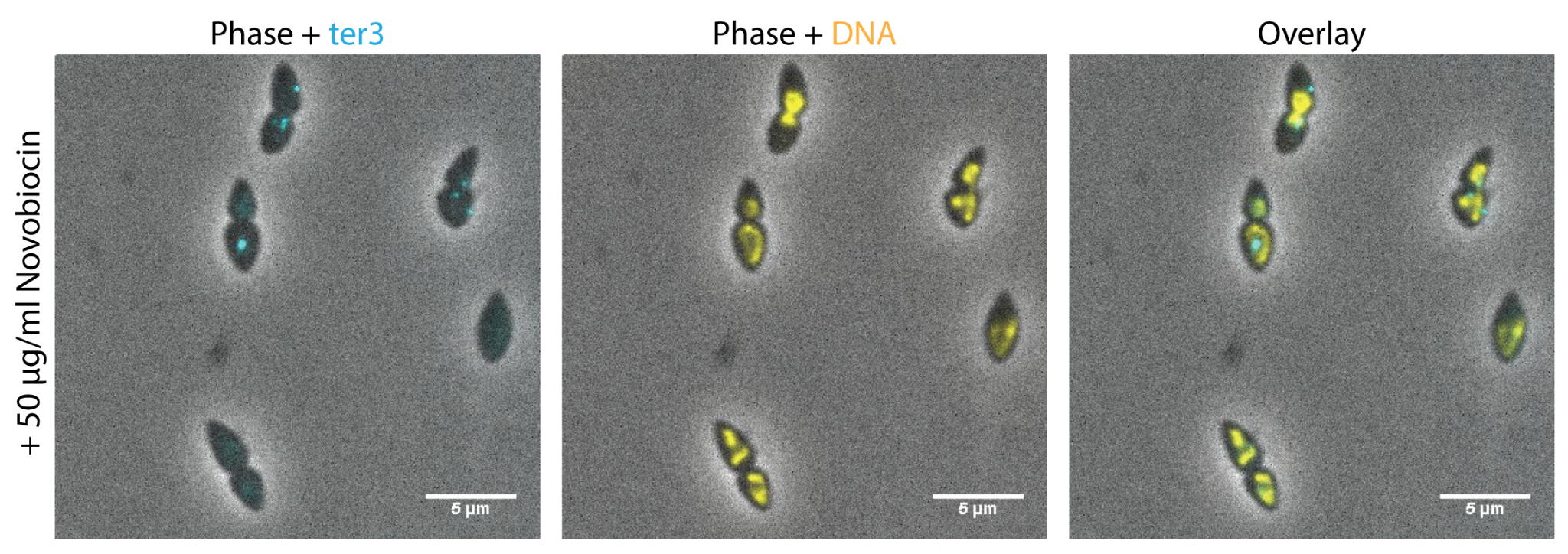

**Figure S6.** Microscopy images of replicating wildtype cells in the presence of Novobiocin (50μg/ml). Phase contrast + ter3 loci (CFP), phase contrast + DNA tagged with HU-mYpet (YFP) and an overlay are shown. Adding Novobiocin had a clear influence on the replication and segregation process in cells. Cells typically had misplaced ter loci with respect to the division septum and often had chromosomes placed in the middle of the cell, rather than at the cell poles. As a result, cells were unable to divide.

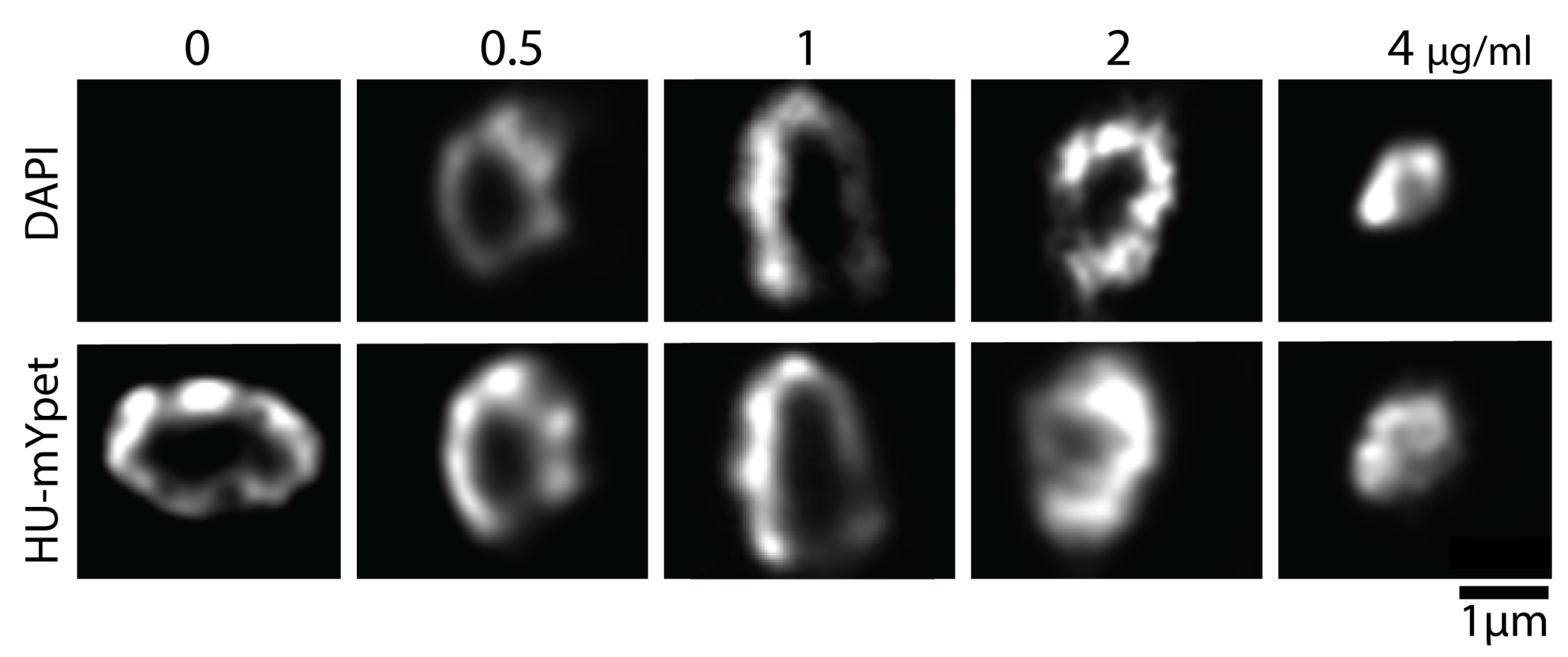

**Figure S7.** Microscopy images of circular chromosomes stained with various concentrations of DAPI and labelled with HU-mYpet. Top: The chromosome is stained with various concentrations of DAPI (0.5, 1, 2 and 4 μg/ml final concentration incubated for 1min). Bottom: The same chromosomes labelled with HUmYPet. High concentrations of DAPI alter the chromosome conformation and compact the nucleoid.
